# Estimating environmental suitability

**DOI:** 10.1101/109041

**Authors:** John M. Drake, Robert L. Richards

## Abstract

**Author statement:** JD proposed the model, JD and RR wrote the code and performed the analysis, JD wrote the first draft of the manuscript, and all authors contributed substantially to revisions.

**Abstract:** Methods for modeling species, distributions in nature are typically evaluated empirically with respect to data from observations of species occurrence and, occasionally, absence at surveyed locations. Such models are relatively “theory-free”. In contrast, theories for explaining species, distributions draw on concepts like *fitness, niche,* and *environmental suitability*. This paper proposes that environmental suitability be defined as the conditional probability of occurrence of a species given the state of the environment at a location. Any quantity that is proportional to this probability is a measure of relative suitability and the support of this probability is the niche. This formulation suggests new methods for presence-background modeling of species distributions that unify statistical methodology with the conceptual framework of niche theory. One method, the plug-and-play approach, is introduced for the first time. Variations on the plug-and-play approach were studied with respect to their numerical performance on 106 species from an exhaustively sampled presence/absence survey of vegetation in the Canton of Vaud, Switzerland. Additionally, we looked at the robustness of these methods to the presence of irrelevant information and sample size. Although irrelevant variables eroded the predictive performance of all methods, these methods were found to be both numerically and statistically robust.

## Introduction

How the occurrence of a species in nature depends on the state of its environment is one of the most fundamental problems in ecology. Conceptual frameworks for answering this question introduce such ideas as *fitness* (Fretwell and Lucas 1969, Hirzel and Le Lay 2008, Peterson et al. 2011), *niche* (Thrasher et al. 1917, Peterson et al. 2011), and *environmental suitabilty* (Engler et al. 2004, Hirzel and Le Lay 2008, VanDerWal et al. 2009, Franklin 2010). Problems with operationalizing these concepts include vagueness about their meaning (Peters 1976, Hurlbert 1981, Orr 2009), failure of real systems to meet assumptions such as *distributional equilibrium* (Elith et al. 2010) and lack of *source-sink dynamics* (Pulliam 2000), and data problems such as the commonness of *presence-only data* (Brotons et al. 2004, Pearce and Boyce 2006, Ward et al. 2009).

This paper seeks to define some of these concepts in a theoretically unifying and computationally operational way. The proposed definitions lend themselves to a new method, which we refer to as the *plug-and-play approach* to modeling environmental suitability. The plug-and-play approach is very flexible. For instance, it allows that data might come from different places at different times. Further, plug-and-play methods for estimating environmental suitability also yield an approach to ecological niche modeling. We study several instances and show that the performance of the plug-and-play approach can be comparable to MaxEnt, a leading method for species distribution modeling (Phillips et al. 2004, Elith, J., Leath-wick 2009, Elith et al. 2011), and superior to a recently proposed density ratio estimator (Kanamori et al. 2009, Sugiyama et al. 2013). Finally, when used in a particular way (*i.e*., using the regularized Gaussian estimator as its base learner), the plug-and-play method can be used for *variable identification* and *fitting of robust models* even in the presence of few records and irrelevant variables.

## Environmental suitability and the ecological niche

The state of the environment at *a* location *i* may be represented by a vector of measurements *z_i_ = [a_i_, b_i_, c_i_,…]* where *a* is rainfall, *b* is temperature, *c* is vegetation, etc. We assume that *z*_*i*_ is constant through time. The *environmental distribution* is the joint density of environments in nature, denoted *f*(*z*) (notation follows Elith et al. (2011)). The *environmental distribution of species s or occurrence distribution* is the joint density of environments in which s is found, denoted *f*_1_*(z)*, distinct from its *range* and *niche* (Drake 2015). The support of *f*_1_, *i.e*., the values of *z* at which *f*_1_ > 0, is the realized niche. The boundary of the support of *f*_1_ is denoted by *h*_*F*_. Sometimes one is interested in the conditional probability that a species occurs at a location given the environment there, *P(y = 1|z)* (Keating and Cherry 2008, Ward et al. 2009, Franklin 2010, Elith et al. 2011, Royle et al. 2012, Hastie and Fithian 2013). We call this conditional probability the *suitability* of environment *z, S(z)*, for species s. By Bayes’ rule,

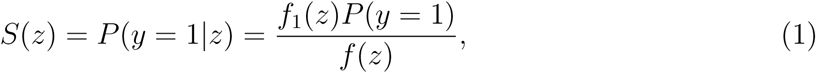

Where P(*y* = 1) is the species *prevalence*. Since prevalence is a proportionality coefficient, we will sometimes wish to ignore it, in which case we have *relative suitability*

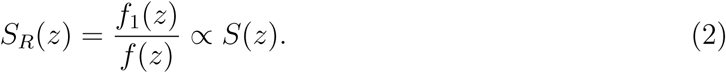

Typically, prevalence will be unknown, although (because it is a single quantity) it might possibly be estimated through independent surveys with less effort than is required to estimate the component densities *f*_1_(*z*) or *f*(*z*) (Phillips et al. 2009). Assuming non-extinction, P(*y* = 1) > 0, *S_R_*(*z*) > 0 if and only if *S*(*z*) > 0. That is, the support of *S* and *S_R_* are identical. The *fundamental niche*, *N*, is defined as the *set of all environments in which the species can persist in the absence of continuous immigration from other populations*. More completely, the fundamental niche, *N*, is the set of all environments *z* such that there exists a population size *n* at which the probability of persisting at a location with environment *z* and in the absence of immigration over a large time horizon *T* ≫ 0 exceeds some threshold θ, possibly close to one, in which case there is a non-zero probabilty that species s will be found in *z* (*S*(*z*) > 0), further implying *S*_R_(*z*) > 0. Typically, this persistence condition will obtain when there is population size n at which average absolute individual fitness exceeds one. This is a cumbersome definition, but it points to a way that such notorious problems as source-sink dynamics (Pulliam 2000), Allee effects (Holt 2009), stochastic extinction (Hanski 1989), and niche conservatism (Wiens et al. 2010) may be conceptually incorporated and quantitatively addressed, rather than having them swept away by unrealistic model assumptions. Formally, 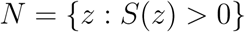. The boundary of this set is designated *h_N_*.

By providing definitions for *environmental suitability* and *niche* and relating these to observable quantities in nature (e.g., the frequency of occurrence of a species), this framework allows us to formulate and answer some fundamental ecological questions, including:

- Is location *i* more or less suitable for species s than location *j* (for any choice of *i* and *j*)?
- What is the dimension of the niche of species *s*?
- What are the environmental variables that comprise the *niche* of species s or influence its distribution?
- How is the potential distribution of species *s* influenced by a given variable *x*?
- What are the boundaries of the niche of species *s*? (What are its environmental *tolerances*?)
- Is the environmental distribution of species *s* set by its niche (its *tolerances*) or by the set of the environments realized in nature?
- How will the environmental distribution of species *s* change when the distribution of environments in nature changes?
- How will the spatial distribution of species *s* change as the distribution of environments in nature changes?

The framework can also be used to empirically address questions in community ecology:

- What is the niche similarity of species *s*_1_ and *s*_2_?
- Do communities *S*_1_ = {*s*_1_, *s*_2_, *s*_3_,… *s*_*n*_}and *S*_2_ = {*s*_*n*+1_, *s*_*n*+2_, *s*_*n*+3_,… *s*_*n*+*m*_} gexhibit more within-community niche variation or between-community niche variation?
- What is the *saturation* or fraction of species in the species pool that could persist in *z*.

## An estimator for environmental suitability

In this section, we propose that this conceptual framework lends itself to modeling, that is building numerical or computer-learned models of environmental suitability or species’ niches from records of species occurrence in nature. Here is the core of the idea. First, many applications do not require an estimate of absolute environmental suitability, but (for instance) only a rank ordering. In such cases, relative suitability is adequate. Particularly, we shall argue below that *niche identification*, which is what we call the process of building a model of *h_N_*, only requires information about relative suitability. The plug-and-play approach to modeling relative environmental suitability proposes that the ratio of two estimates 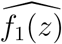 and 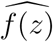, be used as an estimator of *S*_R_:

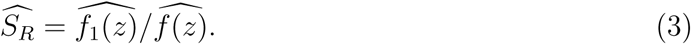

Because *f*_1_(*z*) and *f*(*z*) are just probability densities, they can be estimated using any of a number of techniques for probability density estimation, such as kernel density estimation or Parzen’s window estimator (Jones and Wand 1995). Alternatively, one might substitute a parametric expression, which is basically what is done by MaxEnt, a maximum entropy algorithm commonly used for species distribution modeling (Phillips et al. 2004). Specifically, the MaxEnt algorithm for species distribution modeling stipulates that 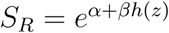, Where α is a normalizing constant chosen so that *f_1_(z)* integrates to one, β is a vector of fit coefficients and *h*(*z*) are transformations of the covariates referred to as *features*. (This function *h* is different from *h_F_* and *h_N_* as defined above, but used here for consistency with Elith et al. (2011).) Thus, MaxEnt is a special case of environmental suitability modeling, but does not adopt the plug-and-play approach, whic*h* allows for the substitution of alternatives for *f*_1_ and *f*, including nonparametric options, as may be suggested by theory (either statistical theory or biological theory), the objectives of a study, or the properties of a data set (such as sample size).

This approach can be extended to the problem of ecological niche identification. For a given estimate of *S*_*R*_, the estimated boundary of the realized niche is just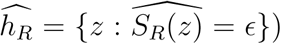, where ϵ > 0 is a small threshold parameter. However, Drake (2015) argued that if a species is rarely found in unsuitable habitats and the environmental distribution is “broad” with respect to the species’ niche (that is, that the range of *f* contains the extreme environments in *N*), then *h*_F_ ≈ *h*_N_ (Drake 2015). Substituting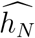 for 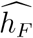, we have an estimator for the boundary of the fundamental *niche*, *i.e.*, 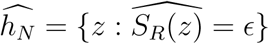.

An alternative to the estimator in equation 3 is to estimate the ratio directly, i.e.,

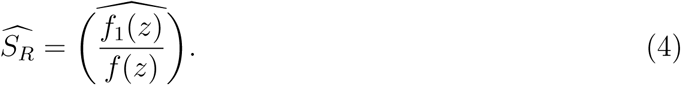

This problem has been addressed generically by Sugiyama et al. (2013) and specifically by Kanamori et al. (2009) in the context of covariate shift adaptation (adapting statistical analyses to changing distributions of independent variables) and Kanamori et al. (2009) and Hido et al. (2011) in the context of outlier detection. The technical similarity between outlier detection and ecological niche modeling has been noted before and used to motivate presence-only models for ecological niche modeling (Drake et al. 2006, Drake and Bossenbroek 2009, Drake 2015). Here the analogy is used to motivate a presence-background approach. For this purpose, Kanamori et al. (2009) introduce unconstrained Least Squares Importance Fitting (uLSIF), which they show to be superior to a variety of alternatives when applied to a simulated classification task. We therefore used uLSIF as a direct density ratio estimator for comparison with the plug-and-play estimator.

## Methods

### Plug-and-play with simple component densities

Implementing the plug-and-play approach requires estimating the component densities *f* and *f*_1_. Candidate estimators for the component density functions include parametric multivariate density estimators (Kotz et al. 2000), robust versions of these (which may be important when data are not normally distributed or contain extreme examples; Huber and Ronchetti (2009)), sparse estimators that reduce variance through “regularization” (which may be important when fitting to small samples from high dimensional spaces; Schäfer et al. (2015)), and nonparametric methods (Jones and Wand 1995).

To compare among these classes, we studied the performance of relative suitability models comprised of three kinds of Gaussian density estimates: (i) ordinary multivariate Gaussian densities, (ii) robustly estimated Gaussian densities, (iii) densities estimated using a shrinkage estimator proposed by Schäfer & Strimmer (Schäfer et al. 2015) in which the pairwise correlation coefficients are scaled by a shrinkage intensity parameter (λ= min(1, max(0,1–λ^*^)), where λ^*^ is the analytic optimal regularization parameter of Ledoit and Wolf (2003)). For comparison, we also studied the nonparametric kernel density estimator of Li and Racine (2003), which would be expected to be superior in cases where one or both of the component densities *f* and *f*_1_ are strongly non-normal, for instance if they are multi-modal or highly skewed.

### Plug-and-play with ensemble component densities

The plug-and-play approach isn’t limited to such simple component densities as elaborated in the previous section, but can be applied to basically any approach to probability density estimation that can be numerically evaluated. Recently, *ensemble methods* have been shown in many areas of statistical application to provide robust probabilistic models that exhibit both low variance and low bias. For instance, random forests are frequently used in classification and regression problems (Breiman 2001). For applications requiring species distribution modeling, the gradient boosting machine is popular (Elith et al. 2008, Elith, J., Leathwick 2009). To be effective, ensemble models must exhibit improvement compared to a single model. Typically this is achieved by “voting” the predictions of a large number of minimally biased (underfit) models (Drake 2014). Ensembles of highly tuned models may even erode performance (Mainali et al. 2015). Thus, it is important when constructing ensemble learners to optimize the entire ensemble, not the base models.

Here we explore how *boostrap aggregation* or *bagging* can be used to improve the estimate of *f* or *f*_1_ by reducing its variance, and thereby improve the performance of the plug-and-play estimator. Bagging is performed by constructing bootstrap samples (with replacement) from a given data set, fitting a model to each, and then reporting the average prediction of ensemble of models (Breiman 1996). Bagging has been found to be a very robust approach to ensemble modeling. Typically, it will be the case that there are many more records of environmental background than species occurrence, so the estimate of *f* will be more precise than that of *f*_1_ (Phillips et al. 2004). Motivated by this observation, we first propose only to bootstrap the estimation of the occurrence distribution in the numerator (*f*_1_), yielding a method we call *NumBag*. In the implementation studied here, a kernel density estimate was obtained for each of *v* = 100 bootstrap samples from the occurrence distribution and averaged before dividing by the kernel density estimate of the background points. Alternatively, we use bagged estimators for both *f*_1_ and *f*, a procedure we call *DoubleBag*. Since the estimated density 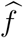 may depend on a very large number of background points, DoubleBag is expected to be much more computationally costly than NumBag. Therefore, it is of interest to determine the performance of NumBag relative to DoubleBag as well as the performance of both in absolute terms. NumBag and DoubleBag are special cases of the plug-and-play idea that offer potentially greater performance than the simple estimators described above, although at a cost of increased computation time.

### Density ratio estimation

In contrast to plug-and-play, the uLSIF algorithm estimates the relative suitability quotient directly. Direct estimation might be desirable if, for instance, estimation of the component densities is more difficult than estimation of the ratio or if estimation errors in the component densities are compounded when the ratio is taken. Sugiyama et al. (2013) discuss density-ratio estimation in general and the uLSIF estimator in particular. One advantage of uLSIF is that it is possible to analytically compute the leave-one-out cross-validation score for a given set of model parameters, greatly decreasing the time needed for model tuning in comparison with numerical cross-validation. MaxEnt (Phillips et al. 2004) may also be viewed as a density-ratio estimator and is probably the most popular approach to species distribution modeling in general.

### Presence-only methods

Finally, these methods were compared to two recently introduced presence-only modeling methods, LOBAG-OC (Drake 2014) and range bagging (Drake 2015). In many cases presence-only algorithms perform as well and in some cases perform better than related presence/absence algorithms (Maher et al. 2014). Briefly, LOBAG-OC applies bootstrap aggregation to low bias (weakly regularized) one-class support vector machines and votes the result. LOBAG-OC, therefore, follows Vapnik’s principle (Vapnik 1998) in that it seeks the set of covariate values for which the probability density is non-zero (the support), rather than attempting to estimate the full probability density (Drake et al. 2006). Range bagging similarly votes a number of base learners trained on bootstrap samples of presence points. In this case, the base learner is the convex hull of a reduced number of environmental covariates randomly selected for each bootstrap sample. The current analysis provides, therefore, not only a comparison of the plug-and-play method with other presence-background estimators, but is also the first comparison of these two presence-only methods themselves.

### Computation

Statistical fitting is straightforward. For this illustration, all fitting was performed in the statistical programming environment R (Alexander Weiße and Fehske 2008). In the case of the ordinary Gaussian density, the covariance was estimated from the unbiased sample covariance using the R function **cov** (Alexander Weiße and Fehske 2008). The robust version was estimated using the minimum covariance determinant (Rousseeuw and Driessen 1999) (function **covRob** in the **robust** package, Wang *et al*. (2014)).The regularized version was estimated using a regularized density estimator (function **cov.shrink** in the **corpcor** package, Schäfer et al. (2015)). Since this estimator truncates the fit distribution to the first two moments, we refer to this as a “regularized Gaussian” estimator. The kernel density estimates were fit using function **npudens** in package **np** (Hayfield and Racine 2008). Bandwidth of this estimator was selected automatically using the analytic rule-of-thumb of Li and Racine (2003). Preliminary experiments suggested that computationally intensive bandwidth selection procedures like cross-validation could improve AUC by an average of only about 0.01 at a cost of ∼ 6000-fold increase in computing time. MaxEnt models were fit using the function **maxent** at the default settings in the R package **dismo** (Hijmans 2012). This function makes use of the java program of Phillips et al. (2004) (available on the web: https://www.cs.princeton.edu/~schapire/maxent/). Density ratio estimates obtained via uLSIF were computed using the R code of Kanamori et al. (2009) (available on the web: http://www.math.cm.is.nagoya-u.ac.jp/~kanamori/software/LSIF/).

### Case study: Ecological niches of sub-alpine vegetation

Drake et al. (2006) studied the performance of support vector machines as niche models using data on 106 plant species in 550 8m×8m plots in the Swiss Alps between 400m and 3200m in elevation. Observed species prevalence (i.e., fraction of sampling plots in which a species was found) ranged from 1.1% to 35.2%. As the above-ground species in these sampling plots were exhaustively enumerated, these data provide a rare opportunity with known absences to benchmark the performance of alternative modeling approaches. Additionally, these data have been studied by Maher et al. (2014), who established that presence-only methods could perform comparably to presence-absence methods and Drake (2014) in an evaluation of the LOBAG-OC algorithm. Together, these papers provide a benchmark against which to compare the performance of the plug-and-play approach to modeling environmental suitability.

Locations were randomly assigned to either training (80%) or testing (20%) subsets. All predictor data were rescaled by subtracting the mean and dividing by standard deviation of samples in the training set. A model was fit for each species using the plug-and-play estimator where 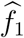 was obtained from the component density fit only to occurrence records in the training set and 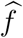 was fit to all records in the training set. Model performance was evaluated by predicting the relative suitability for each record in the test data set and computing AUC, the area under the receiver-operator characteristic, which is a measure of a model’s discriminative ability (Phillips and Elith 2010). Models were tuned using 10-fold cross-validation on the training set with mean AUC calculated for each model on each species across the 10 folds. After fitting, tuned models were evaluated on the test data set. To investigate calibration and the propensity of these models to be overfit by cross-validation, mean AUCs from cross-validation were compared to the test AUC with Spearman rank-order correlation.

### Effect of irrelevant variables

As the volume of automatically recorded and remotely sensed environmental data accelerates, an increasing problem for modeling environmental suitability or ecological niche identification is the determination of relevant variables and optimizing performance in the presence of irrelevant information. Intuitively, one expects model performance to decline with the number of irrelevant variables as the learning algorithm has to sift a smaller and smaller fraction of true correlates from among the many possibilities. Additionally, the number of parameters sharing the available degrees of freedom increases with the number of irrelevant variables, diminishing the amount of information available to estimate each and eventually resulting in ill-posededness for those methods lacking a regularization scheme to solve this problem. On the other hand, it has recently been shown that, amazingly, in some cases pattern recognition is actually improved by the presence of irrelevant variables (Helmbold and Long 2012).

We studied the performance of plug-and-play, LOBAG, range bagging, and MaxEnt methods in the presence of irrelevant variables by simulating 1, 2, 4, 8, 16, or 32 normally distributed random variates with mean zero and unit variance and combining these with the ten genuine variables prior to model fitting. As before, model performance was evaluated by calculating AUC on the withheld test data. Results were summarized by inspecting the erosion of performance (decline in AUC) with respect to the number of irrelevant variables, a measure of statistical robustness, and by tabulating the number of models that converged, a measure of numerical robustness.

### Learning rate analysis

Finally, we studied the effect of sample size on model performance. An important property of any statistical model is the way in which its performance changes with the number of instances available to learn from, the *learning rate*. This problem is particularly acute in species distribution modeling, where a species may be known from only a small number of unique localities. To investigate the learning rate of the algorithms studied here, we identified the 8 plant species with the greatest number of occurrence records in the training data set ( *Geranium sylvaticum, Anthyllis vulneraria, Polygonum viviparum, Achillea millefolium, Lathyrus pratensis, Astrantia major, Plantago media, and Pimpinella major*). Training points were randomly assigned to 10 cross-validation folds. Within folds, each method was trained on randomly selected subsets of the data ranging in size from *n* = 2 to the total number of available presence points in the smallest set of training folds. Each of these models *(m_2_,m_3_,…, m_n_)* was then applied to the points in the test fold and an AUC score for each was calculated as above. These results were visualized by plotting the mean test AUC for each learning method as a function of the training set size.

## Results

Using the classical Gaussian for both component distributions, the plug-and-play method performed moderately well when it could be fit (mean AUC: 0.761), which was approximately 85% of cases (Fig. 1). Models using the robust Gaussian for both components typically yielded a poorer fit (mean AUC: 0.717) and could be fit in a similar fraction of cases (83%; Fig. 1), whereas models using the regularized Gaussian routinely performed very well (mean AUC: 0.829) and could always be fit. The model using kernel density estimates for the components was similarly robust in the sense that a reasonable model could always be obtained. Initially, we thought this improvement might come at considerable cost in terms of computational complexity to allow for empirically optimizing bandwidth parameters, but ultimately we found the model could be estimated using the analytic rule of thumb of Li and Racine (2003) with very little loss of performance (mean AUC: 0.840). The ensemble approaches NumBag and DoubleBag were among the best performing methods we studied, with high average AUC values (0.836 and 0.837, respectively) and the large majority of models able to be fit (98% and 97%). LOBAG-OC (mean AUC: 0.756, 93% fit), range bagging (mean AUC: 0.777), and uLSIF (mean AUC: 0.789) all performed less well. Thus, both the plug-and-play method (when using either KDE or regularized Gaussian component distributions) and the ensemble approaches NumBag and DoubleBag were found to perform comparably to MaxEnt (mean AUC: 0.841; Fig. 1) and better than two other methods we have recently introduced.

**Figure 1:**
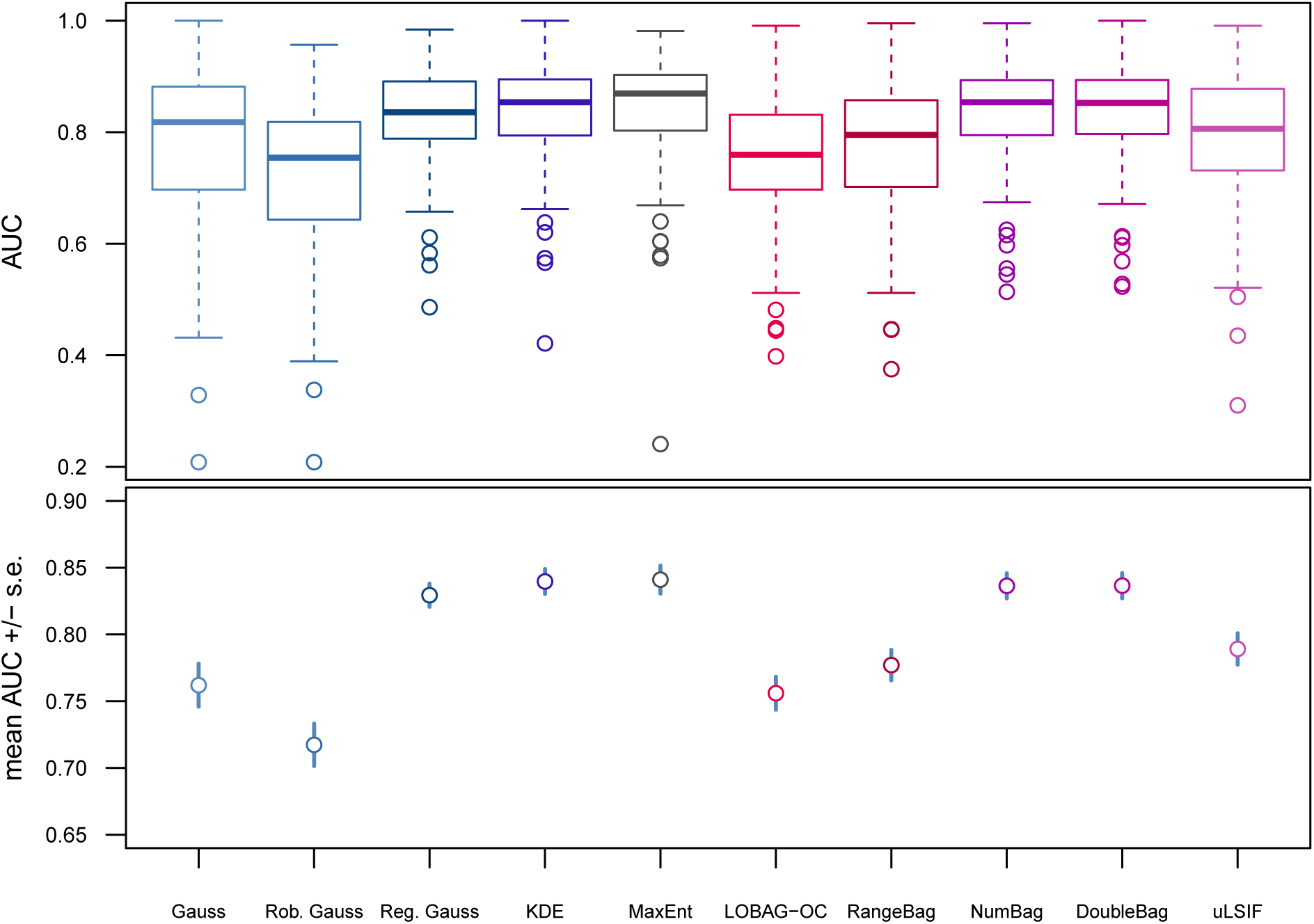
Performance of six plug-and-play species distribution models compared with two presence-only species distribution models (LOBAG-OC and RangeBag), MaxEnt, and a density ratio estimator (uLSIF). The plug-and-play methods were classical multivariate Gaussian densities (Gauss), robust Gaussian (Rob. Gauss), regularized Gaussian (Reg. Gauss), and kernel density estimator (KDE) applied to both *f* and *f*_1_; Numerator Bagged plug-and-play (NumBag); and Double Bagged plug-and-play (DoubleBag). Range in AUC was smallest for the regularized Gaussian, KDE, NumBag, and DoubleBag methods (top panel). Methods with the highest average AUC were the regularized Gaussian, KDE, NumBag, and DoubleBag plug-and-play approaches and MaxEnt (bottom panel). The classical Gaussian and robust Gaussian plug-and-play methods and presence-only methods LOBAG-OC and Range-Bag performed considerly more poorly; the density ratio estimate obtained using uLSIF was intermediate. Error bars are mean +/- s.e. Both the classical Gaussian and robust Gaussian were unable to fit all models due to numerical instability or ill-posedness (89 and 87 out of 106 fit, respectively).

To assess the tunability of each method and vulnerability to overfitting, we compared AUC in the cross-validation folds with the AUC calculated on the test data. An overfit model will have higher AUC on training data than on test data. Additionally, the correlation between the mean AUC in cross-validation folds and test AUC indicates the extent to which the observed AUC in training is predictive of the AUC that can be expected with respect to unseen examples. In this study, all cross-validation AUC values were significantly (*p* < 0:01) and positively correlated with test values (Fig. 2). The strongest correlations were shown by the plug-and-play method with classical Gaussian components (ρ = 0.589), NumBag (ρ = 0.569), DoubleBag (ρ = 0.601) algorithms, suggesting that measured performance will be most indicative of future performance in novel analyses performed with these three models.

**Figure 2:**
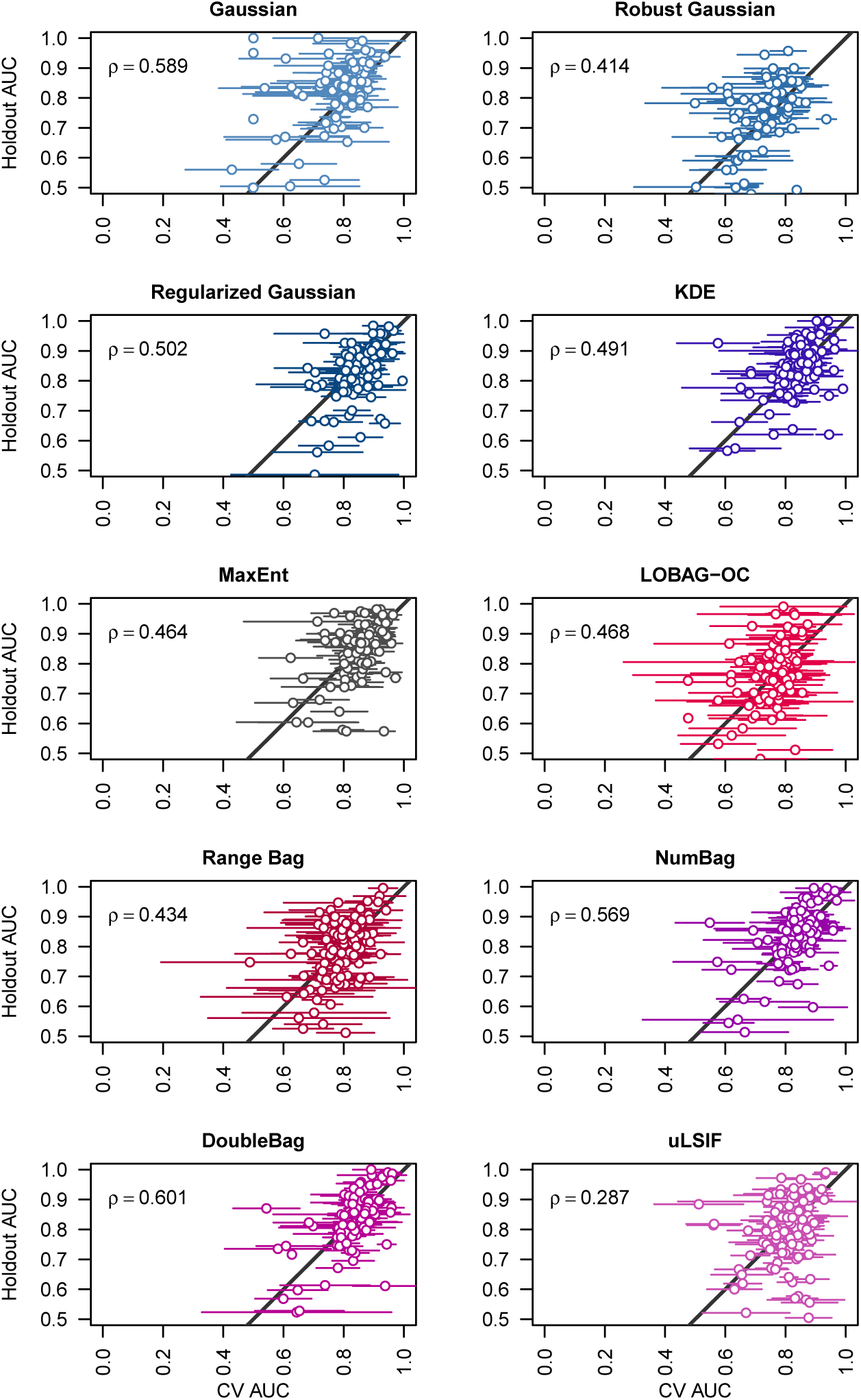
Mean performance (AUC) across 10 cross-validation folds is compared to performance (AUC) of each model on the testing subset. A one-to-one line is plotted on each graph and 95% confidence intervals for mean cross-validation AUCs are shown. Points falling on the one-to-one line are models where the estimated performance in cross-validation was equal to performance on the test set. Points below the line are overfit (higher performance in cross-validation than with withheld data) while points above the line are underfit (higher performance in withheld data than in cross-validation). AUC values were significantly and positively correlated for all models (Spearman’s ρ shown).

Performance of the studied methods varied greatly in the presence of irrelevant variables (Fig. 3). The plug-and-play approach with KDE and regularized Gaussian component densities, NumBag, DoubleBag and MaxEnt models clustered together as the best performing overall (Fig. 3). Specifically, although performance declined as the number of irrelevant variables increased, it declined very slowly so that the reduction in AUC was only about 6% in the presence of 32 irrelevant variables, at which point random variables outnumbered real variables by more than three to one. Thus, in the sense that introducing irrelevant information had little effect on the final output, these three models were all statistically very robust. In contrast, the performance of the plug-and-play approach with classical Gaussian and robust Gaussian component densities, range bagging, LOBAG-OC, and density ratio estimation with uLSIF declined much more rapidly (Fig. 3). Additionally, although the plug-and-play approach with KDE and regularized Gaussian components and MaxEnt models converged in all cases, the plug-and-play approach with classical Gaussian and robust Gaussian components increasingly failed to converge as the dimension of the environmental data increased.

**Figure 3:**
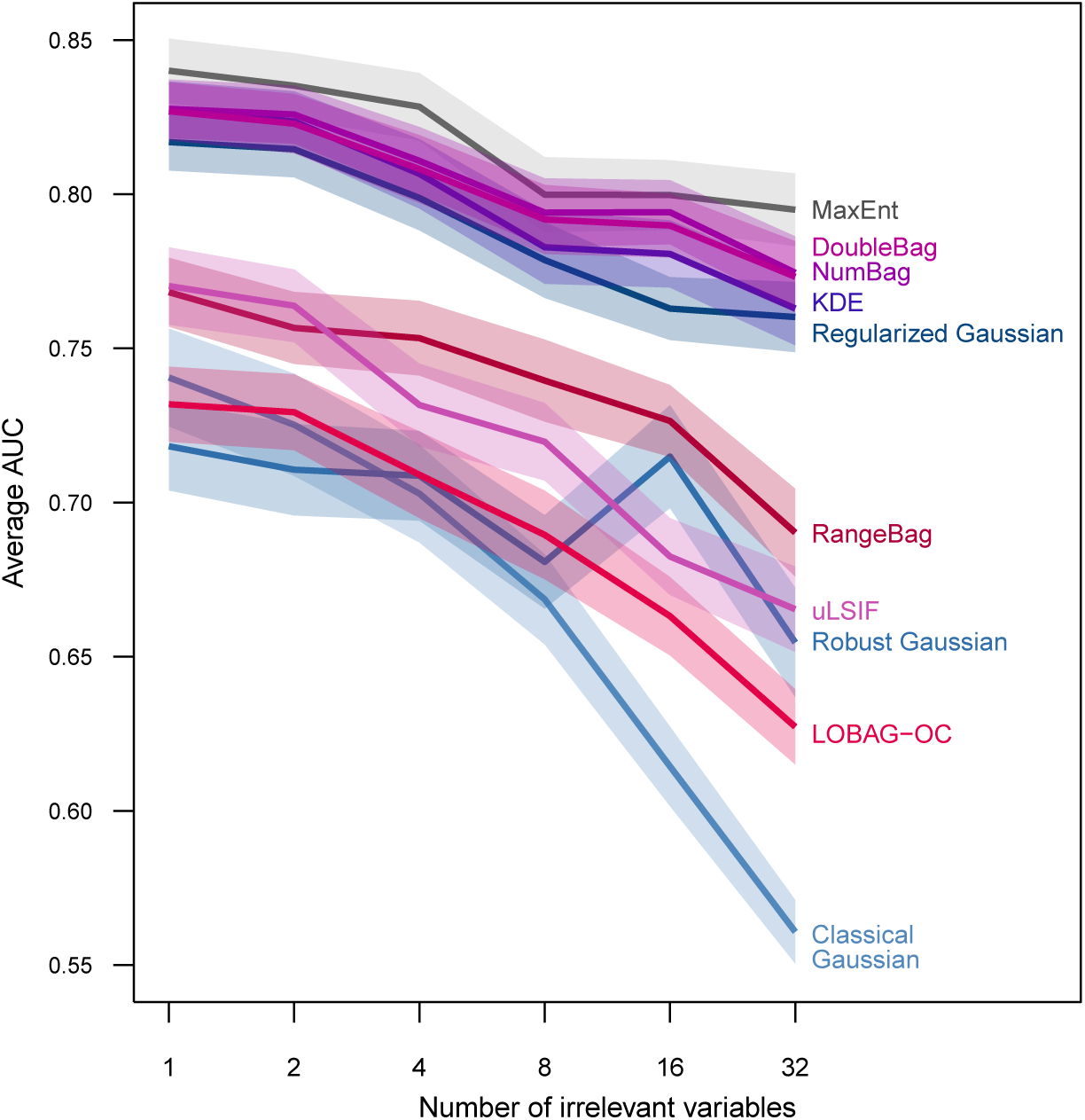
Performance of six plug-and-play species distribution models compared with two presence-only species distribution models (LOBAG-OC and RangeBag), MaxEnt, and a density ratio estimator in the presence of irrelevant variables. Performance was degraded the least by irrelevant variables in the regularized Gaussian, KDE, DoubleBag, NumBag, and MaxEnt models (top panel; error bars are mean +/- s.e.).

Analyses of learning rate showed similar performance for some methods across the eight most abundant species in our data and variable performance for others (Fig. 4). Additionally, the niches of some species (*e.g.*, *Plantago media*) seemed inherently more learnable. The group of high-performing methods (*i.e.*, DoubleBag, NumBag, KDE, regularized Gaussian, and MaxEnt) tended to perform relatively poorly with fewer than around 15 training points, but improved quickly approaching their maximum performance at around 20 or 30 observations. At the other end of the spectrum (*i.e..* with around 100 observations in *Achillea millefolium and Plantago media*), MaxEnt tended to perform slightly better than DoubleBag and other high performing methods, although this was not always true (compare *Pimpinella major*). Interestingly, the performance of the density ratio estimated with uLSIF varied substantially across species. In some cases (*e.g*., *Pimpinella major* and *Plantago media*) uLSIF performed consistently less well than the group of highest fit models, while in other cases (*e.g.*, *Geranium sylvaticum* and *Lathyrun pratensis*) uLSIF overtook the performance of DoubleBag with increasing numbers of training points. The robust Gaussian method was a clear outlier overall, routinely exhibiting lower performance at all sample sizes.

**Figure 4:**
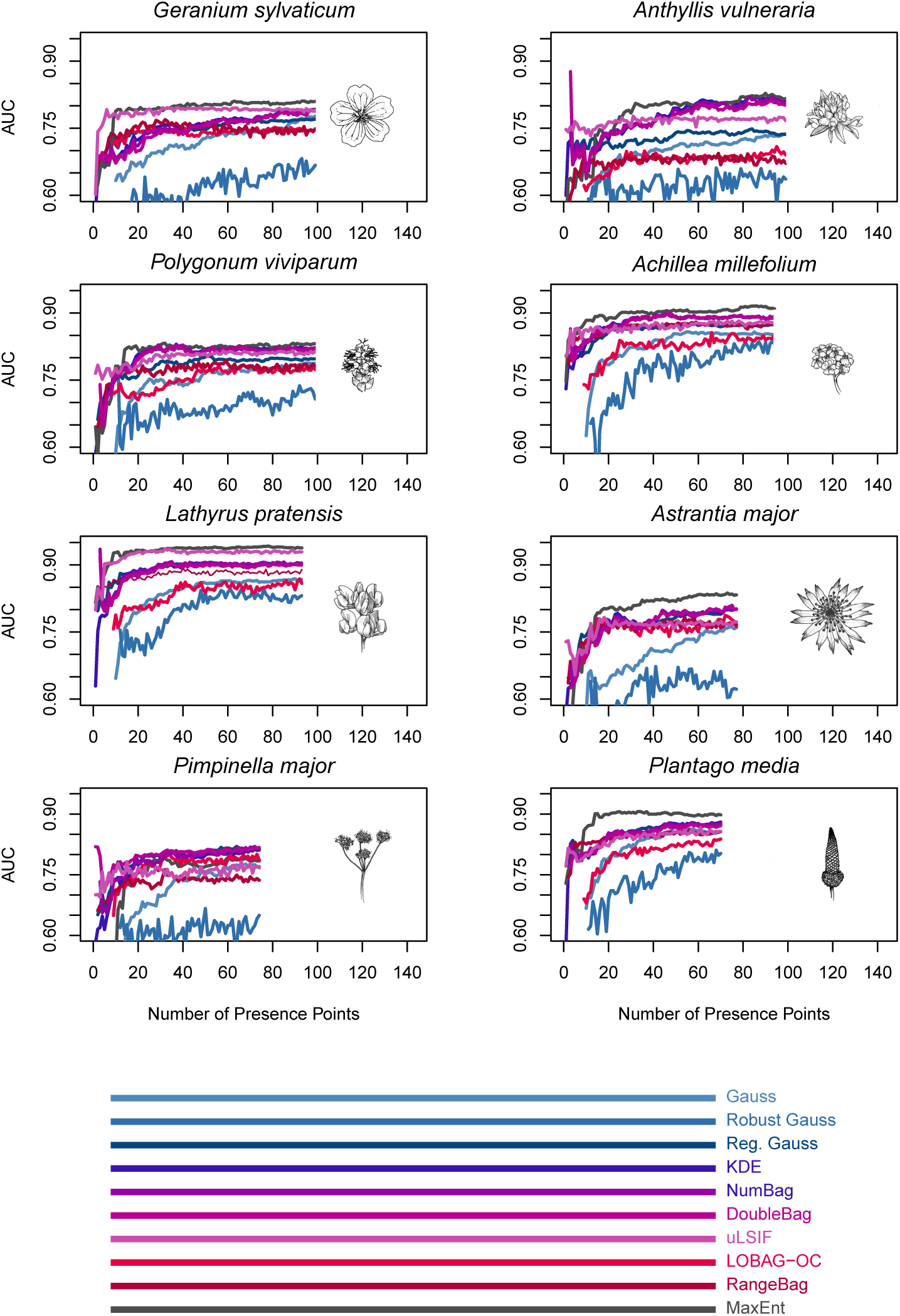
Learning rates of six plug-and-play models, two presence-only models, MaxEnt, and a density ratio estimator on data for 8 abundant species. Models generally perform more poorly at low numbers of training points and gain performance as more training points are oered. KDE, NumBag, DoubleBag, and regularized Gaussian tended to approach a performance plateu at large samples sizes while uLSIF shows inconsistent behavior, occasionally outperforming all other models at all sample sizes *(e.g., Geranium sylvaticum)*, but more commonly showing superior performance only at small sample sizes *(e.g., Anthyllis vulneraria)*.

## Discussion

This study proposes numerical definitions for the ecological concepts of *environmental suitability, relative suitability*, and *niche* that we believe are consistent with common usage. A virtue of these definitions is the potential to unify ecological niche theory with dynamical theory, as alluded to by Peterson et al. (2011), particularly where those theories make probabilistic statements about occurrence within a landscape (*i.e.* metapopulation theories) or persistence over long times (*i.e.* stochastic population theory). A practically useful feature of this framework is that it suggests a new approach – the plug-and-play method – for estimating relative suitability.

Performance of six versions of the plug-and-play method was studied with respect to 106 plant species that had been exhaustively sampled for species presence and absence. Mean AUC of these models ranged from 0.717 to 0.841. Except for the plug-and-play approach with classical and robust Gaussian components, the current models are all superior to those of Drake et al. (2006), which used support vector machines in three different ways and achieved average AUC of less than 0.8 in all cases. On the other hand, Drake et al. (2006) fit models only to data on species occurrences (*i.e.*, presence-only models) whereas plug-and-play is a presence-background approach. Similarly, Maher et al. (2014) studied nine presence-only and seven presence/absence niche modeling methods using these data. All sixteen of those methods returned average AUC of less than 0.8, with the exception of a k-nearest neighbor method configured to fit both presence and absence data where the average AUC was 0.804. Thus, the presence-background approaches studied here (with the exception of classical and robust Gaussian components), are superior to all the presence-only and presence-absence methods studied by Maher et al. (2014) with AUC improvements on the order of 4-6%. One conclusion from this study together with earlier analyses (Drake et al. 2006, Drake 2014, Maher et al. 2014) is that there may yet be scope for improvement, but also that many methods may be contrived to yield very similar results. Particularly, a number of methods perform similarly to MaxEnt, which is currently a widely preferred method. Ultimately, if ecologists wish to choose a method on the basis of empirical superiority (compare Brotons et al. (2004), Elith et al. (2006)), many more data sets will be needed.

One of the key assumptions of the approaches introduced here is that environments are constant and species distributions do not primarily reflect past environmental conditions. Modeling environmental suitability in the presence of *dynamic environments* is an important problem for further research. Extensions of the current approach to dynamic environments are easy to imagine. Suppose we supplement our spatial indexing (*i*) with a temporal indexing (*t*). Occurrence records also must be indexed by time. Recognizing that a location’s past environment affects its present species composition, one would model the probability of occurrence as a function of both present and past environments, perhaps downweighting the the effect of conditions in the distant past or considering only those environmental conditions within a window of time. Now, the probability of occurrence is given by a function of both present and past environments or a weighted mixture of densities. The definition of environmental suitability with respect to the present time is retained, however. A simpler way to implement this idea is to introduce time-lagged covariates into the feature set, *i.e*., *z_i,t_ = [a_i,(t)_,a_i,(t−1)_,a_i,(t−2)_,…b_i,(t)_, b_i,(t−1)_, b_i,(t−2)_,…]*. Joint densities *f* and *f*_1_ could be fit as done here and present suitability returned either by marginalizing over historical environments or by evaluating the full model (including time-lagged variables) for all locations’ actual environmental histories. As the current study of irrelevant variables shows, if the lagged environmental variables aren’t grossly irrelevant or too numerous, the effects they have on a suitably chosen model (*i.e*., plug-and-play approach with KDE or regularized Gaussian components or MaxEnt) are minimal, and we may expect the resulting model to be fairly robust.

In principle, it might seem that this approach also makes the assumption that the species distribution is at equilibrium (since it is assumed that *f* and *f*_1_ are stationary). Applied naively to all the data that one might acquire, this is true. However, because the problem does not explicitly include spatial dependence, the modeler can easily restrict the data to a subset prior to estimating *f* and *f*_1_. Thus, for instance, the set of records for estimating *f* and *f*_1_ could be just the locations that were searched for the species. (In fact, this is what has been done in any study where the background data reflect only study locations, including the current study.) Alternatively, the data could be selected to be drawn only from regions considered to be “accessible” to the species (Peterson et al. 2011). This is very similar to the problem of study extent addressed by Barve et al. (2011).

This paper advocates estimating relative suitability or niche boundary rather than absolute suitability in applications where species’ prevalence is irrelevant (for instance to rank sites by their value to species conservation). The plug-and-play approach nevertheless solves an unnecessary intermediate problem: estimating the densities *f* and *f_1_* (or, in the case of sample selection bias, f_2_). It would seem that estimating the ratio *f_1_/f* directly might be a more efficient approach, as has been advocated by Sugiyama et al. (2013). In comparative studies, Sugiyama et al. found models based on density ratio estimation to be superior to models based on density estimation itself (Sugiyama et al. 2013). The empirical analysis reported here shows this not to be generally true, however, as uLSIF exhibited performance inferior to plug-and-play methods. Except for MaxEnt (Elith et al. 2011), to our knowledge, density ratio estimation has not previously been used for species distribution modeling. What, then, explains the superior performance of MaxEnt compared with uLSIF? Possibly it is due to the fitting criterion, which MaxEnt stipulates to be the Kullback-Leibler divergence between *f*̂_1_ and *f*̂. In contrast, plug-and-play methods propose no such constraint. Thus, for instance, in the current study the alternative component distributions are chosen to meet different criteria applying to different plausible conditions that might restrict the performance of a simple estimator (like the classical Guassian). As it turns out, some of these options degrade performance (i.e., robust Gaussian components), while others improve it (*i.e*., KDE and regularized Gaussian components). The set of possible estimators for the component densities *f*_1_ and *f* and the rationale for choosing among them is an important problem for further study. For now, we recommend the plug-and-play approach with KDE or regularized Gaussian components or MaxEnt when computing resources are limited. In contrast, when the size of the data is small or when computing resources are not a concern, NumBag and DoubleBag are our method of choice.

## Acknowledgments

This work was begun in 2012 while JMD was a Leverhulme Visiting Professor at Oxford University. Additional support was provided by the National Oceanic and Atmospheric Administration Agreement No. NA10NOS4780218.

